# The life of P.I. Transitions to Independence in Academia

**DOI:** 10.1101/571935

**Authors:** Sophie E. Acton, Andrew Bell, Christopher P. Toseland, Alison Twelvetrees

**Author notes:** **Correspondence:** | | |.

## Abstract

The data in this report summarises the responses gathered from 365 principal investigators of academic laboratories, who started their independent positions in the UK within the last 6 years up to 2018. We find that too many new investigators express frustration and poor optimism for the future. These data also reveal, that many of these individuals lack the support required to make a successful transition to independence and that simple measures could be put in place by both funders and universities in order to better support these early career researchers. We use these data to make both recommendations of good practice and for changes to policies that would make significant improvements to those currently finding independence challenging. We find that some new investigators face significant obstacles when building momentum and hiring a research team. In particular, access to PhD students. We also find some important areas such as starting salaries where significant gender differences persist, which cannot be explained by seniority. Our data also underlines the importance of support networks, within and outside the department, and the positive influence of good mentorship through this difficult career stage.

## Introduction

Academic careers have expanded across the university sector in the past couple of decades at all career levels – from post-graduates to professors. However, this is a pyramidal career structure consisting of very few levels: PhD, Post-Doc and group leader. PhD and postdoctoral training positions typically offer time-limited positions therefore the only route to continue in academia is to become an independent group leader. This leads to a highly competitive and tough system. We know this first-hand since we are all newly independent ourselves, starting our own positions 2016/2017 at universities in the UK. We designed a survey for our peers, for all new group leaders, principal investigators, and new lecturers to try and understand the variation in how we are recruited, how we are supported, and what criteria we are judged against as we get established as independent group leaders. Whenever there is a fierce competition, it is important to make sure fairness applies across the sector.

There is no single or linear route to become an independent group leader in the UK. The most frequent routes are recruitment as a permanent lecturer or as a fixed-term research fellow. The latter could be funded directly by universities or through external sources such as the research councils and larger charities. Lecturers typically follow a probation scheme which leads to confirmation of their appointment whilst the situation for research fellows is more precarious. However, externally-funded research fellows typically have significant funding available for long-term research projects and positions to recruit lab members, allowing more rapid establishment of research projects compared to lecturers. Within the 4 weeks that this survey was open we heard from 365 newly-independent researchers in the UK. These were predominantly from the life sciences (83%) but also included physical sciences, social sciences and clinician scientists. Now we use these data to understand what works well but also where support is missing. We find that our peers comprise a resilient and determined group and that some simple measures could make a large impact in supporting our early stages of academic research careers.

## Results

Our cohort consisted of 365 respondents which represents a significant proportion of new group leaders in the UK recruited over the past 6 years (**Figure 1**). A majority are from the life sciences. While this may represent a limitation of the cohort, it may also be a true reflection on the availability of new positions within life sciences – there are more fellowship opportunities in the life sciences.

**Fig. 1.**
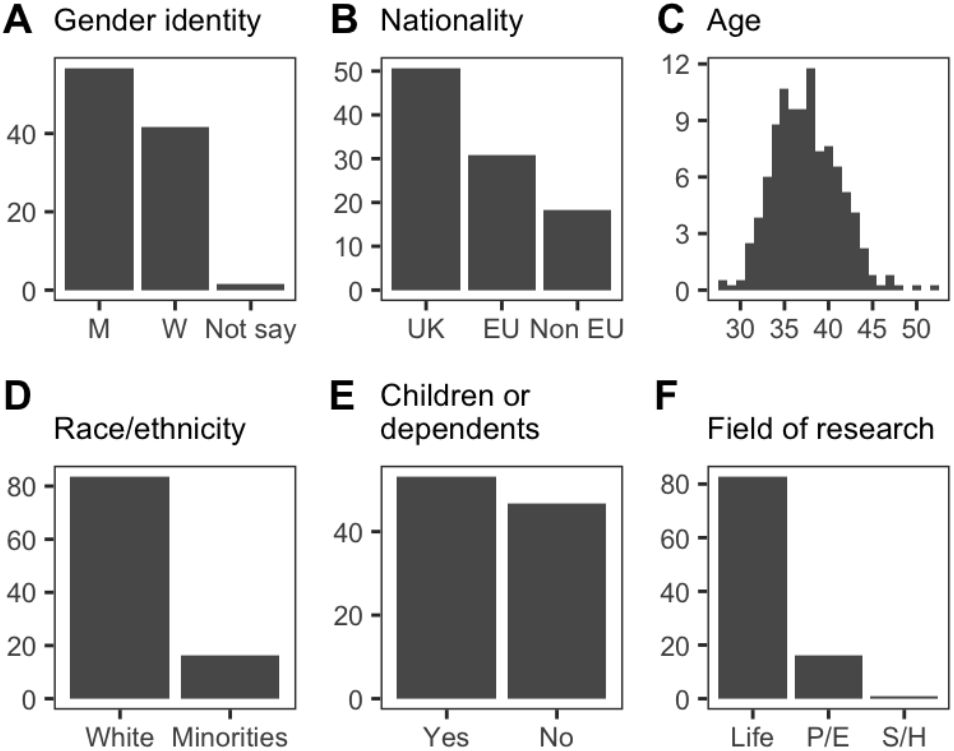
Overview of cohort demographics. All plots are expressed as the percentage of respondents. **A**, Categories are Men (M); Women (W); and ‘Prefer not to say’ (Not say). **B**, Nationality of participants was grouped into those from the UK, other EU countries (EU) and the rest of the world (Non EU). **C**, Mean Age of participants at the time of the survey was 37. **D**, 83.6% of respondents were white. **E**, 53.2% care for dependents.

In general, our cohort was in their mid-thirties and approximately half looked after dependants (**Figure 1C,1E**). As expected, our cohort consisted of more male than female researchers (57% male) (**Figure 1A**). Latest statistics have shown continued disparity in the numbers of male to female professors (approximately 70% male) within academia (www.advance-he.ac.uk), despite good gender equality in the numbers of postgraduate trainees and postdoctoral researchers. Our data show that we are still not at 50:50 in the recruitment of new group leaders, but it is a promising sign for a better balance in the future. 50% are not from the UK, with 30% coming from the European Union (**Figure 1B**). This statistic represents the international mix within academia, the appeal of the UK but also the potential loss of talent through brexit, and lastly the overall importance of international mobility within the sector. Over 80% of our respondents classified themselves as white (**Figure 1D**), which although seems high, is in fact in line with the national average for the UK. There has been some discussion about a lack of role models in academia for minority students, and while this may be true currently, this dataset shows that the next generation of professors may do some good in redressing the balance.

The average researcher spent 7 years between their PhD and starting their own group (**Figure 2A**). Based upon the typical funding period for a post-doctoral position, researchers would have 2-3 fixed-term positions before gaining independence. Those postdocs that have made the successful transition to an independent position will probably have 1st author publications from each of their postdoctoral positions, which is quite an achievement in these short fixed term contracted positions. The 7 year post-doc period reflected in this data set is likely determined by the eligibility restrictions (years post PhD) for career development fellowships that have been in place with most funders, until very recently. It will be interesting to observe whether this changes significantly following the recent decision by research councils and many charities to remove time limits for the fellowship schemes. A longer time spent as a postdoctoral researcher may allow some individuals the extra time needed to publish ambitious or collaborative projects which require longer timelines. On the other hand, extending this time frame may push the age of newly independent researchers higher than currently (average at 34 years in this survey). This means that postdocs are more likely to be balancing starting families, while on short-term contracts with pressure to show mobility. International mobility plays a key role in the academic career path. 51% of respondents had spent >1-year training outside of the UK as postdocs. 67% of respondents had undertaken at least one international move as part of their career (either when moving between PhD and Postdoc or moving from Postdoc to independence in the UK) (**Figure 3**). With both brexit and an anti-immigration political atmosphere, it will be important to put in place visa systems for academics to permit continued international movement.

**Fig. 2.**
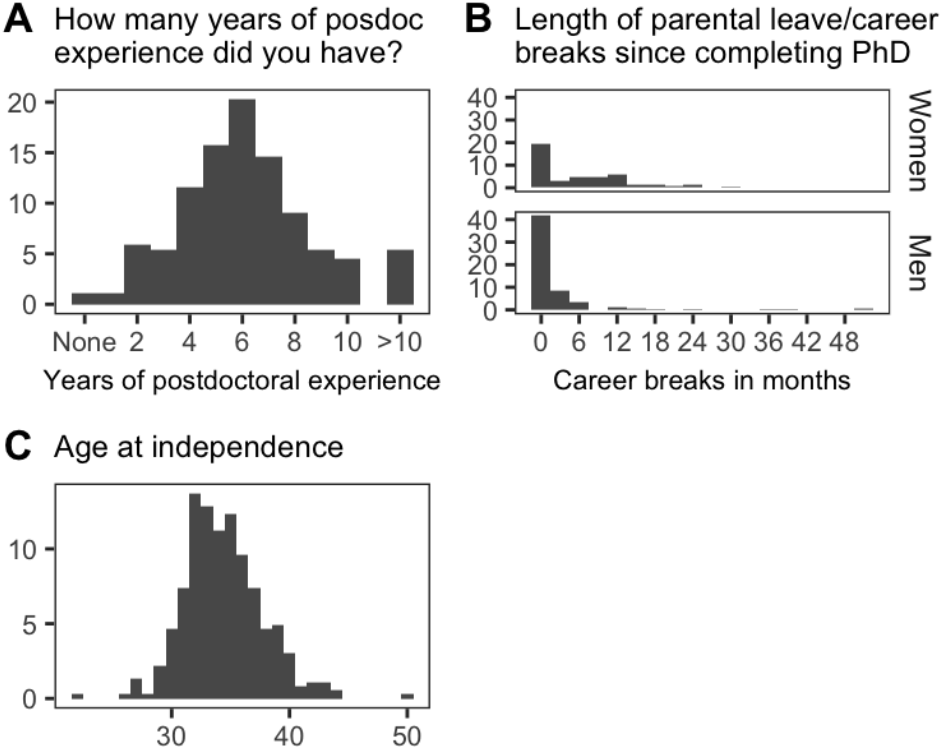
Postdoc experience, career breaks and age at independence. All plots are expressed as the percentage of respondents. **A**, 49% of respondents had between 6 and 8 years postdoc experience prior to independence. **C**, Mean age at independence was 34.

**Fig. 3.**
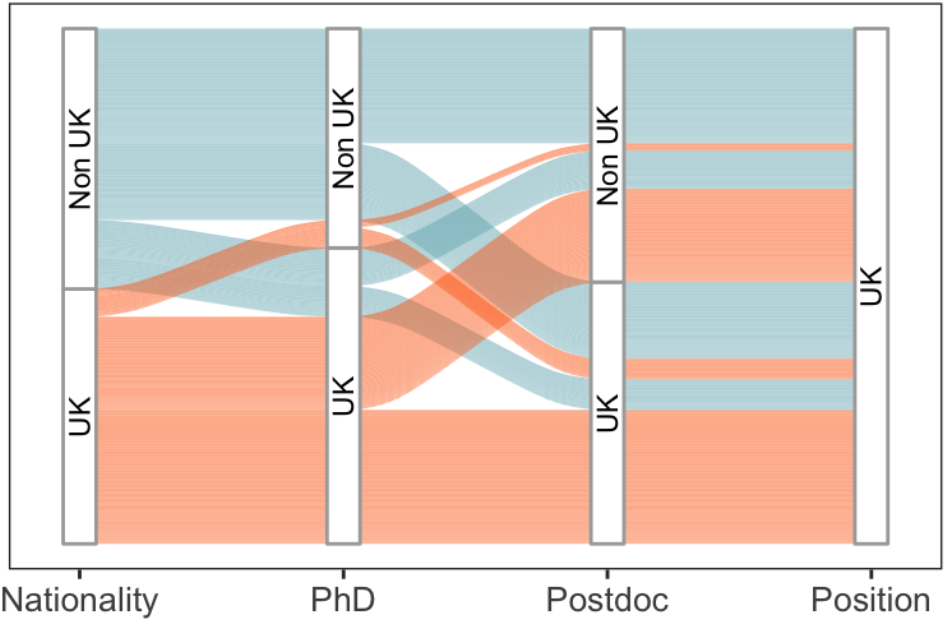
Overview of international mobility through career progression.

### Job Satisfaction and well being

While details about our cohort reveal information about the sector as a whole, it is when we look at job satisfaction of our cohort that we can begin to reveal where problems may lie for new group leaders. More than 50% of new PIs were satisfied with internal factors such as their departments, host institutions and space/facilities (**Figure 4**). However, that leaves at least a third that did not express satisfaction and approximately 20% were dissatisfied or very dissatisfied. We want to use the data collected here to highlight the factors behind these problems and suggest what support could be put in place for these individuals.

**Fig. 4.**
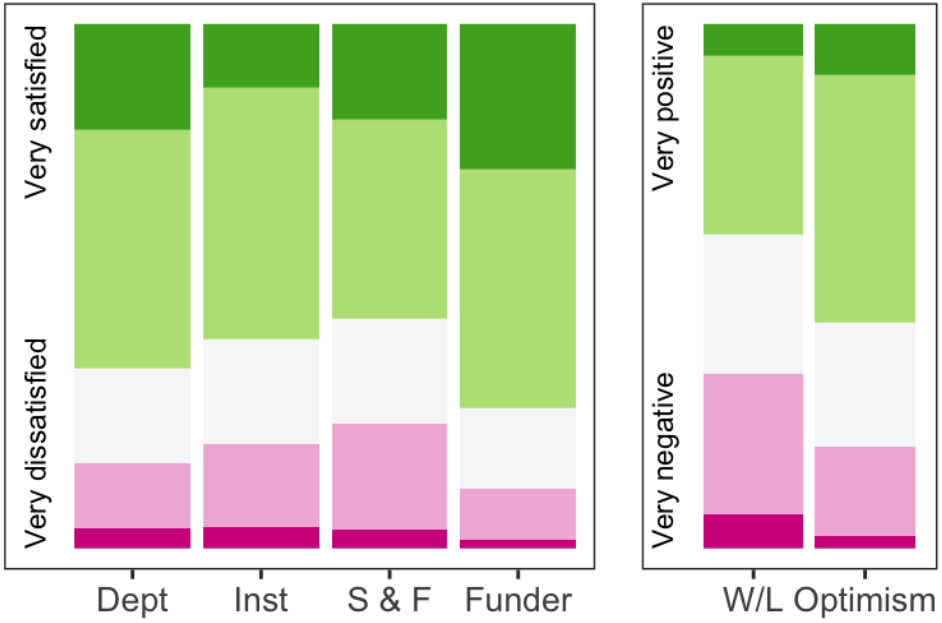
Satisfaction and optimism. Participants were asked to rate their satisfaction with their host department (Dept), host institution (Inst), lab space and access to facilities (S & F) and support from their funder (Funder). Participants were also asked how they felt about their current work/life balance (W/L) and their optimism about their future career (Optimism).

Overall, respondents were least satisfied with the space and/or facilities (**Figure 4**). Lack of space and facilities may be a situation funders can try to improve through communication with host institutions. This will be discussed more at the end of this report. The funders themselves carry the highest level of satisfaction, despite the highly competitive nature of gaining funding (**Figure 4**), but presumably many of our satisfied respondents were those who had successfully navigated applying and winning these competitive grants and fellowships. Work-life balance carried the largest dissatisfaction with 34% dissatisfied. Work-life balance difficulties and increasing time pressures are frequently reported in academia, and almost none of our respondents were working part-time (**Figure 5A**). While flexible working is available through most employers, it appears this option is incompatible with starting a research group. This is not surprising considering the different strains put upon new academics: find funding, build research group, prepare and give lectures and publish work; all within limited time scales and while in competition with other groups internationally. These factors are better expressed in direct quotes from our respondent:

> “I feel like I’m trying to do three separate jobs (research, management/admin, teaching) as well as be a mother… be my own postdoc (because I can’t afford one), be the lab technician (because I can’t afford one), be the lab manager (because I can’t afford one…), be a good mentor for my students, plan strategy, write grants (constantly, I need the money), stay up to date with other research, prepare new teaching material (this takes me ages, I want to do a good job), teach, mark assessments and answer student queries etc. I could go on. No, seriously, is it even possible?”
>
> “I’m confused as to the direction I should be taking and what is really expected of me”
>
> “finally obtained security of a permanent contract but currently feel like crippling teaching load has all but ended my research career.”

**Fig. 5.**
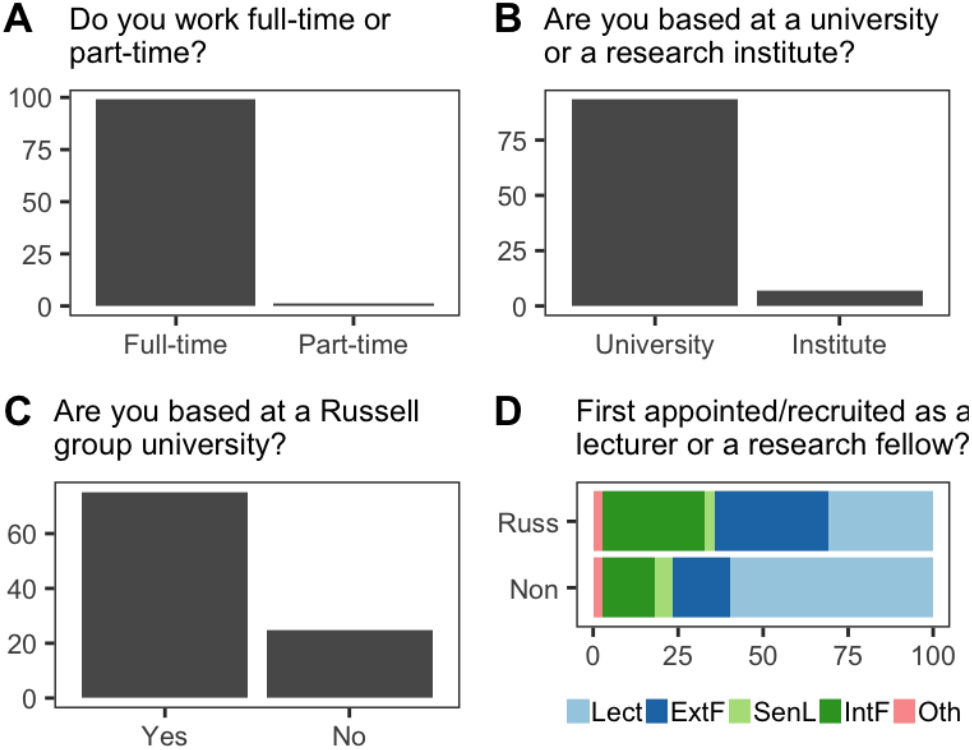
Working patterns, host institution and initial recruitment. Plots **A-C** are expressed as the percentage of respondents, **D** is expressed as the percentage of respondents within each category. Abbreviations: Non, Non-Russell Group University; Russ, Russell Group University; Lect, Lecturer; ExtF, External Fellow; InF, Internal Fellow; SenL, Senior Lecturer; Oth, Other.

Clearly there is a need to find other mechanisms to improve the work-life balance besides offering part-time/flexible working patterns. However, despite elements of dissatisfaction, it is important to highlight over 50% of respondents were optimistic for the future (**Figure 4**). This highlights a strong resilience and positivity amongst new group leaders as they tackle the various demands of their role. It was also encouraging to find that having dependants did not affect satisfaction ratings or optimism scores for either male or female PIs. That we as junior PIs are now able to balance work and family/childcare commitments, and that is has become normal for both mothers and fathers to use some of the flexibility that academic careers provide to achieve this balance, is very reassuring, and suggests that there has been significant change in culture and support for working parents in recent years. Female investigators had unsurprisingly taken more career breaks/parental leave (**Figure 2B**), but it is encouraging to see that fathers are also taking significant periods of leave and sharing childcare responsibilities.

### Career track comparison and gender disparity

As previously mentioned, there is no single route to independence in the UK academic system. Therefore, we wish to compare the individuals within two of these routes (**Figure 5**): Lecturers versus research fellows. Approximately an equal number of research fellows and lecturers were captured in the survey. The research fellows secured funding from a range of bodies but 70% of the respondents were from Russell group universities (**Figure 5C**). This puts a large amount of resources into these 24 institutions. We do not have the data to determine how this is spread across the UK, but funders should look to check what proportion of funding is being awarded to the so-called ‘golden triangle’ of Oxford-London-Cambridge. 65% applied for an advertised position (lecturer or internal fellowship) and the majority of people (>75%) changed department or institution as they transitioned to independence (**Figure 6**). This has become standard practice is recent years and obviously would cause a large degree of disruption and logistical issues for spouses and dependants. There is a lot of pressure to show mobility through PhD and postdoctoral training contracts, and there are major benefits to doing so, however as the average age for starting an independent research group increases, and may only increase further, this becomes an issue for many.

**Fig. 6.**
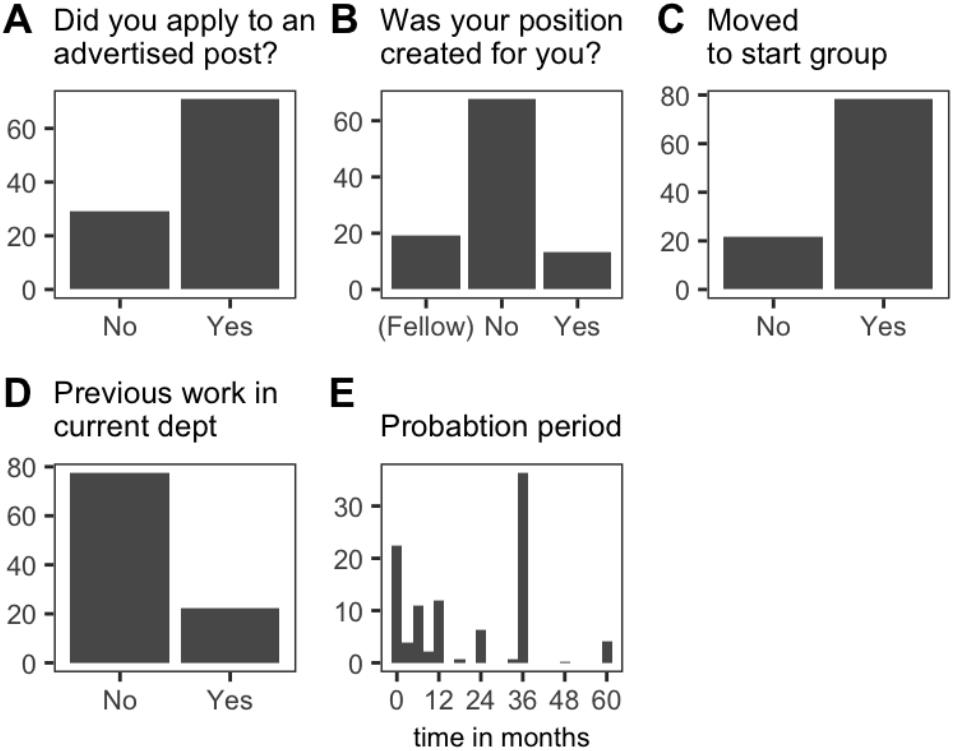
Recruitment of new principal investigators. All plots are expressed as the percentage of respondents

**Fig. 7.**
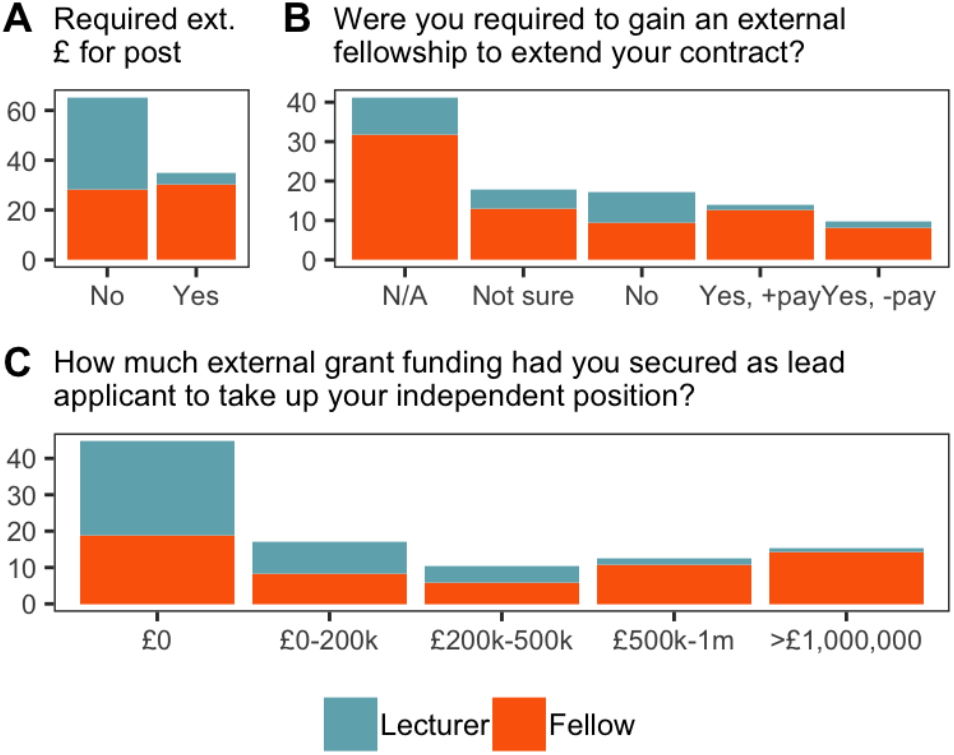
Comparison of lecturers and research fellows, recruitment and external grant funding. All plots are expressed as the percentage of respondents. Abbreviations: “Yes +pay”, Yes including my salary; “Yes-pay”, Yes, but host pays my salary.

38% of our PIs were required to successfully apply for major grants or fellowships in order to take up their position (**Figure 7**), with research fellows bringing in the highest levels of funding with 25% securing >500,000k before starting their position (**Figure 7C**). Despite this large income generated for their host institution, only 30% of research fellows had a proleptic appointment during their fellowship (**Figure 12**), which is discussed further below.

We start to see gender disparity within grant funding very early in a PI’s career, despite what should be an equal starting point at this career stage. Female PI’s are already lagging behind their male peers when we measure whether our respondents had secured additional funding (**Figure 8A-C**). The majority of our cohort (male and female) had secured some additional funding within the first 5 years, but the additional funding won by female PI’s was significantly lower in overall value than their male counterparts, with men much more likely to have secured additional funding worth >1million (p=0.025), and women had also been awarded significantly fewer grants (p=0.039) (**Figure 8B-C**). It looks as though male new investigators were better able to gain momentum and accelerate through continued grant success, allowing them to build critical mass expanding the numbers in their labs, whereas female investigators were more likely to stall and 5 years into running their research group were often failing to gain momentum with funding and therefore recruitment. This is also reflected in Figure 15, where there is a trend for fewer PhD students, fewer postdoctoral fellows and overall smaller lab size when the PI is female (**Figure 15**). This is a worrying trend and we do not have the data in this report to understand why this is the case. We might want to consider however that when it comes to promotions and senior fellowship or programme grant applications, the female PIs will be lagging behind their male counterparts 5-6 years in. We should delve deeper into this issue and ensure that female PIs are being encouraged and supported to apply for more funding and to build their teams in the same way as male new PIs.

**Fig. 8.**
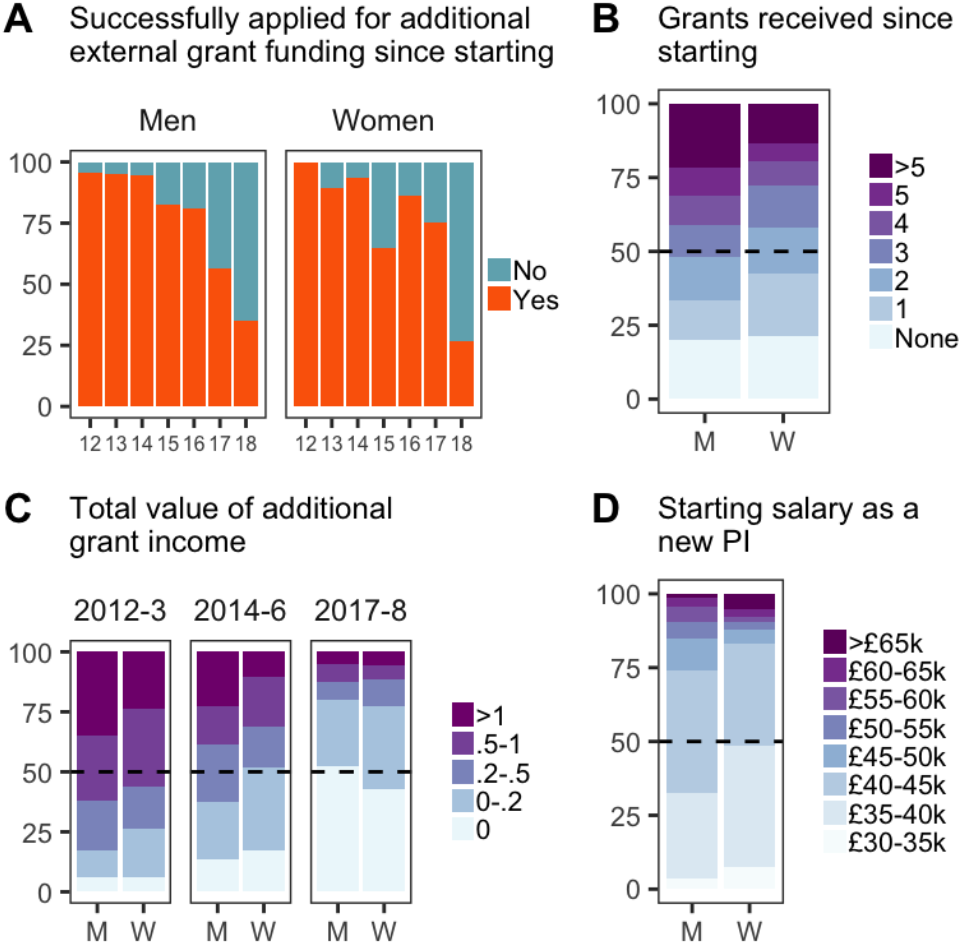
Gender comparisons. Abbreviations: M, Men; W, Women. **A**, Grant success versus year of independence. **B**, Half the men had received three or more grants since starting, half the women received two or more. **C**, Grant values expressed in millions of pounds over three new investigator cohorts based on starting year. **D**, Self reported salaries of new investigators at the time they were appointed show a substantial gender pay gap. All plots are expressed as the percentage of respondents within each category.

Research fellows start on higher salaries than lecturers, but the research fellow salary is not funded by the university. This finding should not come as a surprise considering the selection process to gain the fellowship, prestige of the fellowship, and that gaining significant funding is a promotion criteria at many universities. Indeed, many fellowship schemes include a salary enhancement for the fellow with this in mind. Conversely, lecturers have permanent positions subject to a probation period typically lasting 3 years and criteria which typically includes specific number of outputs, funding success and various levels of teaching. However, even when accounting for the difference in salary between the different career tracks, we found in this data set that the majority of female PIs where being paid less than their male counterparts (**Figure 8D**). This corresponded to a 3-5k annual difference but more crucially was determined by which grade their appointment was made at; grade 7 vs grade 8 (lecturer) and grade 8 vs grade 9 (senior lecturer). It seems that through negotiations with host intuitions, female applicants are more likely to be appointed at the lower of two possible grades, and although this makes little difference to actual salary initially, it has huge implications for future career progression and promotion. It would be useful to collect data on the success rate and timing of promotions of PIs to professorships, to understand if there is a problem here.

Somewhat surprisingly, the majority of research fellows (57%) were still expected to teach as part of their role despite their salary being paid by their funder rather than the university (**Figure 9A**). Although the number of contact hours was significantly less than those in lectureships, nearly 40% of research fellows were expected to contribute more than 10 hours of lectures, or tutorials to undergraduates (**Figure 9B**). Some research fellows, 10-15%, were taking on >40 hours of lectures(**Figure 9B**). Essentially, research funders are directly paying for some proportion of this university teaching. There is an argument that fellows should engage with their departments, bring new material to undergraduate courses, and participate in some level of teaching early in their independent careers, and indeed if the new PI is aiming to be appointed as a lecturer in the longer term then engaging with teaching early should be beneficial for their career development. However >40 hours of lectures and tutorials seems excessive for a new research fellow and all funders might need to consider specifying a limit on the number of hours to protect their investment in the research programme. Some funders do already specify a limit in teaching hours for fellows, but these data suggest that this limit may not always be being enforced by the fellow or respected by the host institution. This wide range in teaching load inevitably impacts the time available for research projects and the likelihood of the career development fellow to successfully apply for a senior research fellowship in the future. Teaching load therefore needs to be part of negotiations with a host institution during the recruitment process and should also be fully transparent to the funders. We would suggest that the best way for newly independent researchers to engage with undergraduate teaching would be to focus on laboratory supervision of undergraduate laboratory projects or literature projects, so that the teaching is directly contributing to their research programme if possible. New PIs should also be aware that supervision of master’s students and PhD students would also count as teaching contribution.

**Fig. 9.**
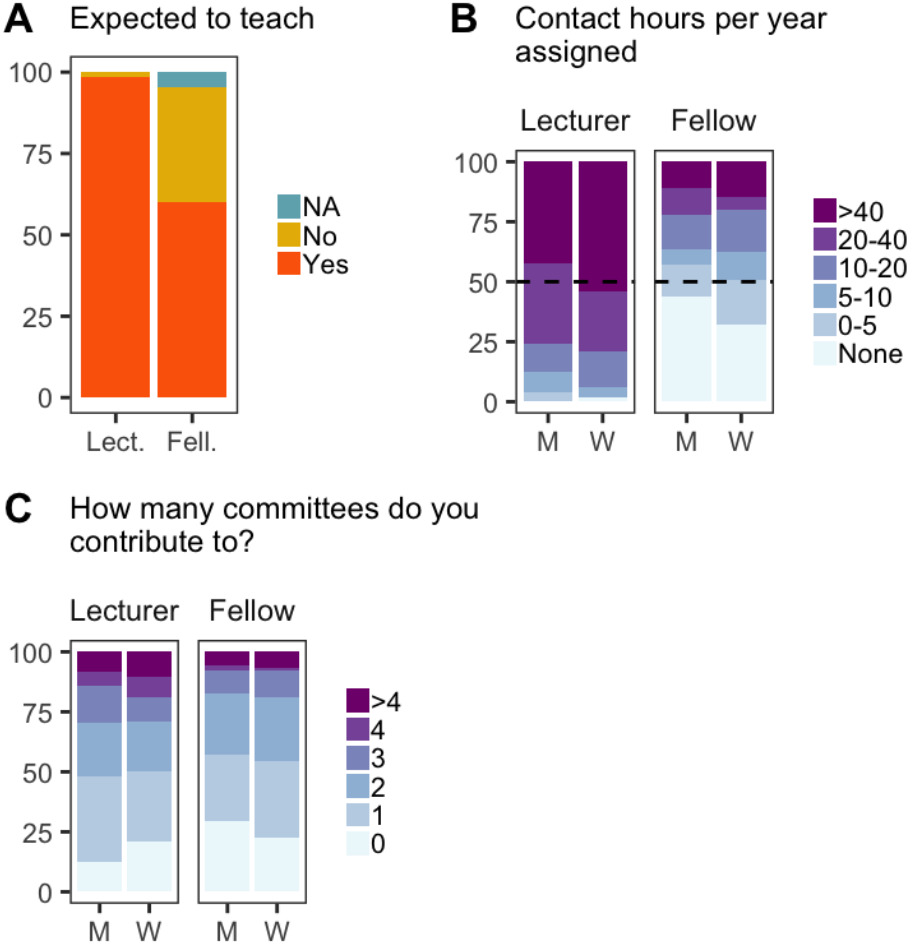
Teaching and administration load. Abbreviations: Lect, Lecturers; Fell, Research Fellows; M, Men; W, Women. All plots are expressed as the percentage of respondents within each category.

Support in a new independent position is especially important as we take on roles we have no prior experience of. We found that almost 25% of all new PIs felt that they had no mentorship (**Figure 10**), and when correlated with the earlier data on optimism, the unmentored women in our data set were the least optimistic for their career progression (**Figure 11**). Most new PIs had an annual review, but research fellows were more likely to be missing out on this advice (**Figure 9C**). Research fellows were also less likely to be members of their university union than lecturers (**Figure 9D**). Nearly 18% of externally funded fellows reported not having an annual review compared with just 3% of lecturers (**Figure 9B**). However, research funders provided an additional source of support for fellows through external fellows meetings, a valuable source of peer support and career advice which was not available to most lecturers, even when in receipt of research grants from major funders (**Figure 9E**).

**Fig. 10.**
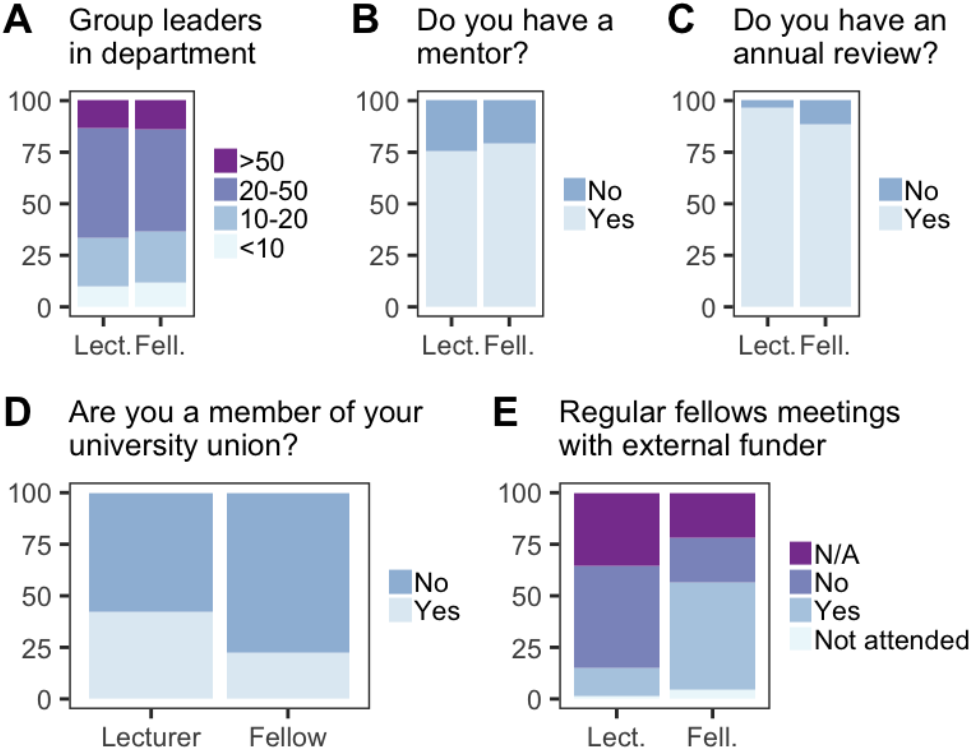
Support and mentorship. Abbreviations: Lect, Lecturers; Fell, Research Fellows. All categories are expressed as the percentage of respondents within each category.

**Fig. 11.**
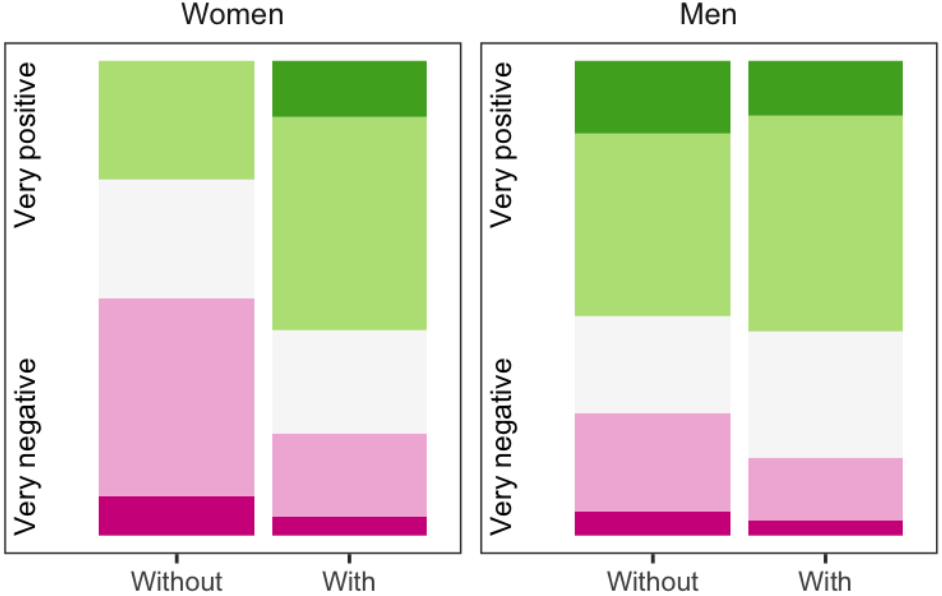
Future optimism between groups with and without mentorship.

The majority of research fellows had no proleptic appointment, or underwritten contract, with the university as they started their position (**Figure 12**). Moreover, 58% of research fellows report they are unaware of clear criteria in order to successfully gain a proleptic appointment or a permanent position (**Figure 13**), and furthermore are not aware of when any formal review process or assessment will be undertaken by their host institution or department (**Figure 13**). This leaves junior PIs in a stressful position without job security, but also without any clear aims. Research fellows support 100% of their salary, they bring in large grants, these individuals have been through stringent external selection processes, and on the whole are more likely to contribute to the REF and the research status of the institution. Therefore, the lack of clear career progression criteria needs to be addressed within the sector (**Figure 13**). The funders should lead this change, to protect their investment in these junior researchers. Many of our respondents reported examples of other junior PIs being let go at the end of their career development fellowships, despite publishing well and taking on teaching responsibilities. Since fellows are hired by universities on contracts which are dependent on the funding source, they can be made redundant at the end of the fellowship with little consequence. Our respondents had a lot to say on this particular matter - many comments extremely critical.

> “Career progression is very non-transparent. Vague descriptions of the areas in which excellence is required, but no idea of the level equivalent to excellence. Getting a proleptic appointment is very difficult”
>
> “It is widely believed that if you have funded your own salary from grants for 7 years then the school should take you on as a full-time lecturer. However this does not appear to be written down anywhere and may have been inconsistently applied.”

**Fig. 12.**
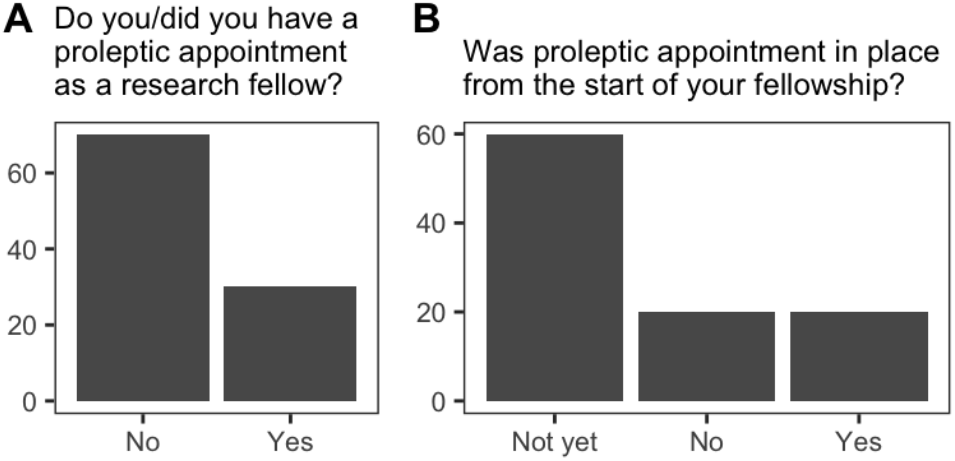
Proleptic appointments for research fellows. Both plots are expressed as the percentage of respondents

**Fig. 13.**
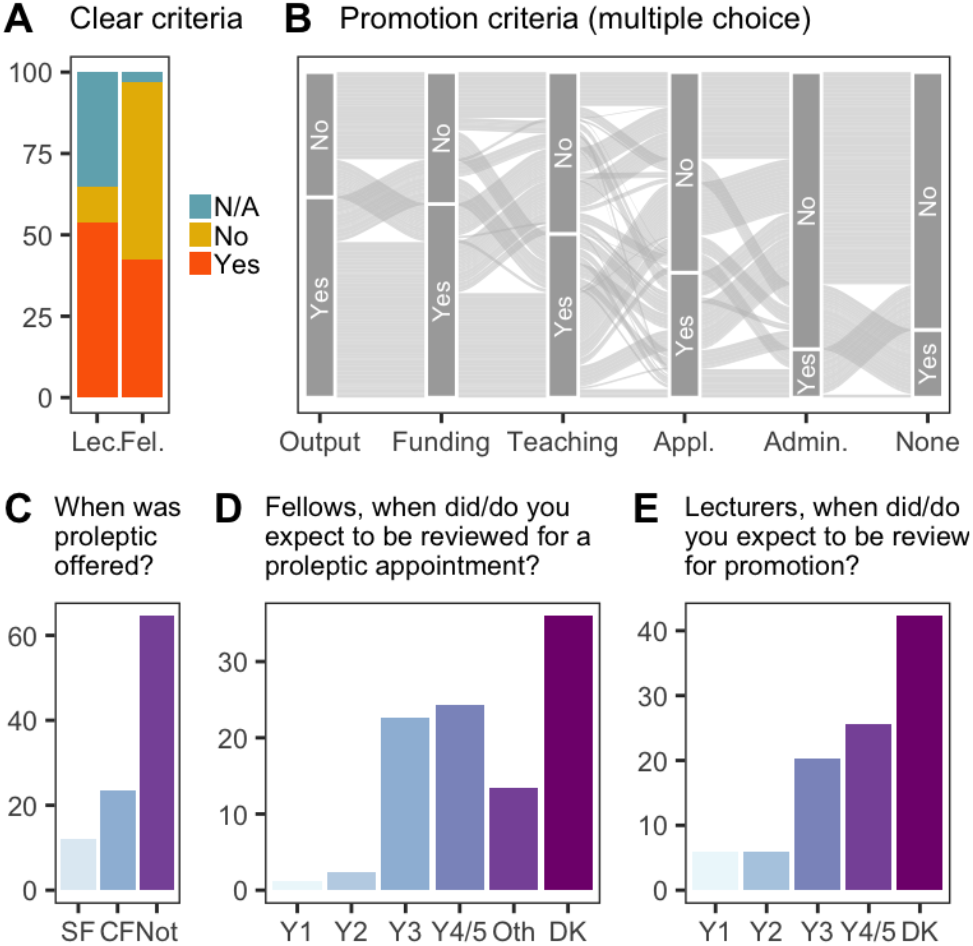
Promotion and probation criteria. Participants were asked if they had clear criteria for probation or proleptic appointment as lecturers (Lec.) or fellows (Fel.) **A**, followed by whether these criteria included **B**: specific outputs (Output); funding success (Funding); teaching load (Teaching); funding applications (Appl.); administrative roles (Admin.); or none of the above (None). Fellows were asked when a proleptic appointment was offered (**C**, after securing senior fellowship, SF; at the end of their career development fellowship, CF; or not reviewed yet, Not). Fellows (**C**) and Lecturers (**D**) were then asked in what year did/will review happen (Year 1 (Y1) to 4 (Y4); other, Oth; Don’t Know, DK). **A-B** are expressed as the percentage of respondents within each category. C-E are expressed as the percentage of all respondents.

With regard to financial support from the university, to build a research group lecturers were more likely than research fellows to be provided with a PhD student by their department (59% vs 41%) (**Figure 14A**), and most lecturers were provided with start-up funds (88%), in the region of £20-60k (**Figure 14F**). Externally funded research fellows were not provided start-up through their department but they are frequently covered via their fellowship. Overall, there was very little financial contribution to research fellows by the university, with most costs covered by their external funders.

**Fig. 14.**
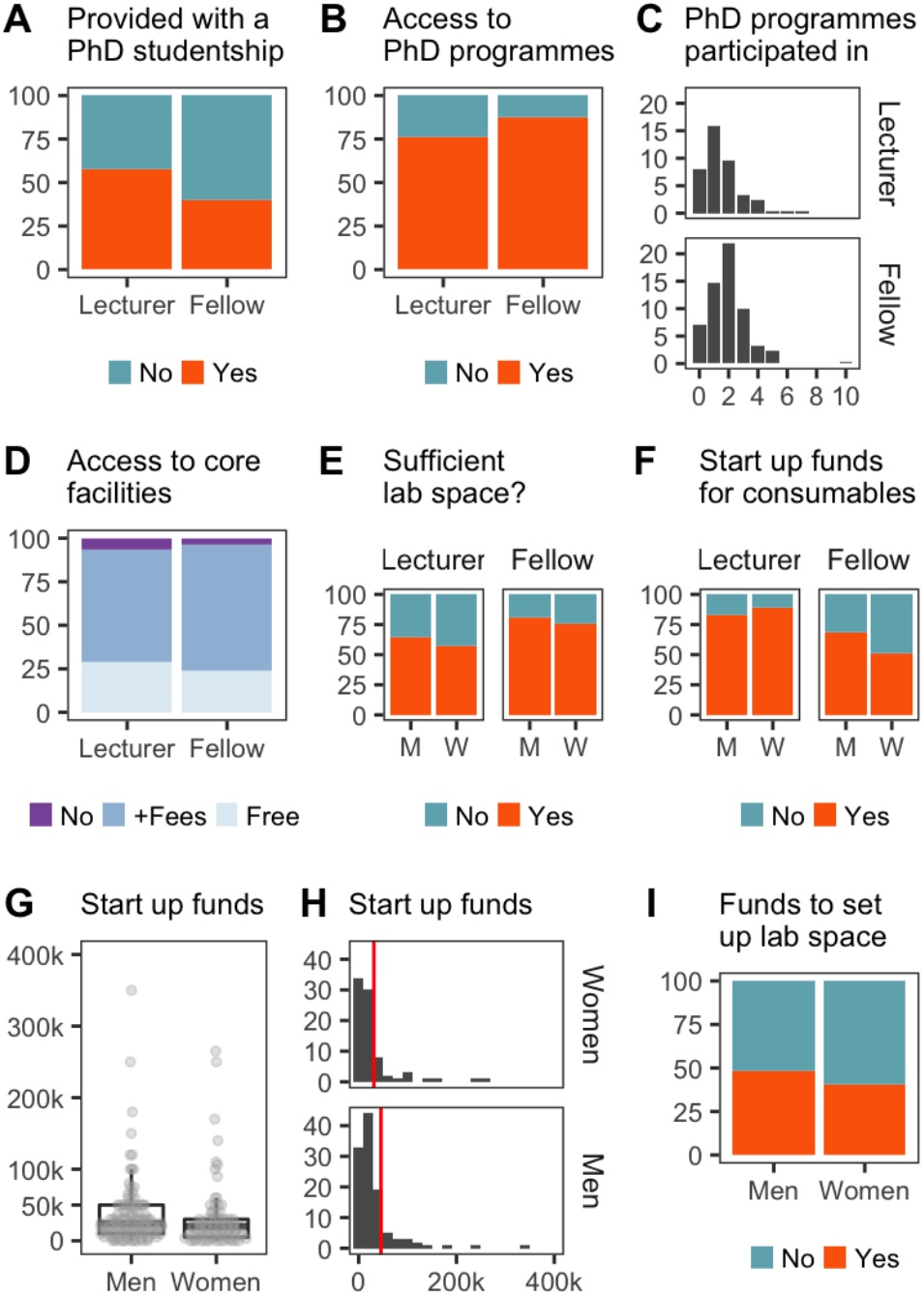
Start up funds, lab space, and PhD programmes. Plots **A-F & I** are expressed as the percentage of respondents within each category. Participants were also asked the amount of start up funds they were provided with (**G-H**). Red vertical line indicates the mean start upfundsfor each category (**H**). Abbreviations: M, men; W, women.

Most lecturers and fellows had access to PhD programmes (**Figure 14B**), and participated by submitting projects to up to 5-6 in some cases (**Figure 14C**). Despite this however, both new lecturers and research fellows reported finding it extremely hard to get access to and recruit PhD students. On the whole, very few funding options exist for individual PhD studentships within the UK. Many schemes have now been gathered into large doctoral training centers. These are typically spread across Russell group universities and labs compete for the positions. More established and senior labs benefit from these schemes, and new labs struggle to both get their project accepted into the scheme and then to persuade students to join their team. Since most career development fellowships fund only 5 years of research, attracting a PhD student in the first or second year is critical for that PhD student to have time to make a significant contribution to the lab within the tight funding time frame, ahead of a senior fellowship application.

> “To be successful as a Fellow it is primordial to get a PhD student during the first year of contract. Without hands in the lab we cannot work. This is not granted, I struggled to get my lab members. Actually I secured a external studentship, but incredibly and annoyingly my Institution does not allow me to be primary supervisor!”
>
> “I was told in no uncertain terms that the department could offer me nothing as a start-up. I am part of 2 possible PhD schemes in the university but funding only has been awarded to senior colleagues.”

50% of lecturers had no postdoctoral researcher in their group (**Figure 15**), whereas 80% of research fellows had at least one postdoc working with them and some had 3 or more within their first 5 years (**Figure 15C**). As expected, research fellows are better positioned to get up and running faster than lecturers, since at least one position is likely funded via their fellowship. Research fellows were also far more likely to have a research assistant in the group (53%), compared to only 25% of lecturers (**Figure 15D**). Pressure, particularly on the lecturer’s time and budgets is also increased by high numbers of undergraduate students in the lab, in the absence of postdocs or research assistants to help train and supervise (**Figure 15E**). If we return to the job satisfaction data, breaking down the answers reveals lecturers were the least satisfied with work-life balance compared to research fellows. Based on the findings above, this is likely to be related to increased pressure on new lecturers to gain research funding and build a research team. As highlighted above, research fellows begin their position with funding and additional research positions and therefore the research activity can begin immediately. Moreover, while research fellows contribute to teaching, their hours are less than lecturers.

**Fig. 15.**
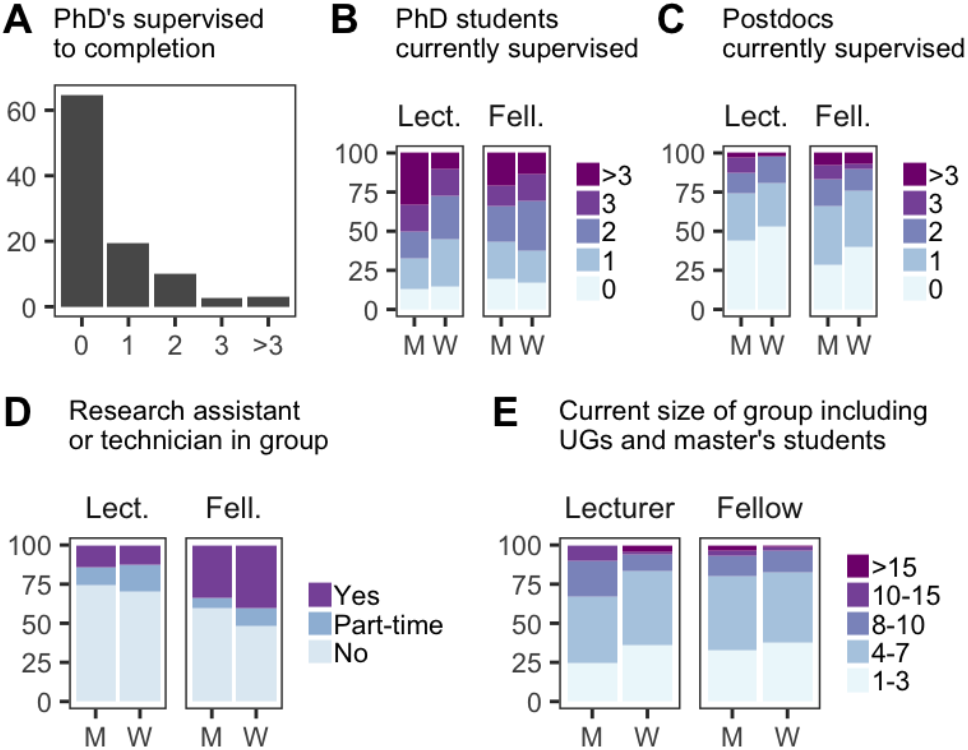
Building a research group. All categories are expressed as the percentage of respondents. Abbreviations: M, men; W, women; Lect., lecturer; Fell., Fellow.

And finally, while we have already described the male-female distribution of the cohort (which is close to a 60:40 split) our data highlighted a very worrying statistic on gender disparity in recruitment around the REF2014 (**Figure 16**). It is perhaps understandable that the lead up to REF2014 stimulated an increase in recruitment of research fellows and new lecturers, however it is extremely clear from our data that this wave of recruitment significantly, if not entirely, favoured male applicants (**Figure 16**). We find this unacceptable and warrants further investigation to understand why the disparity is so extreme in these circumstances. We suggest that this wave of male recruitment may have been due to an increase in direct head-hunting, or more informal recruitment techniques driven by networks within fields, and that these practices seem to favour men. Knowing that this happened in 2014 gives us a chance to highlight the issue, and to hopefully push host institutions to be more mindful with their recruitment ahead of the the next REF cycle. The rules on whether outputs are transferable for submission to the REF have also changed this round, which may remove this wave of recruitment the year before submission.

**Fig. 16.**
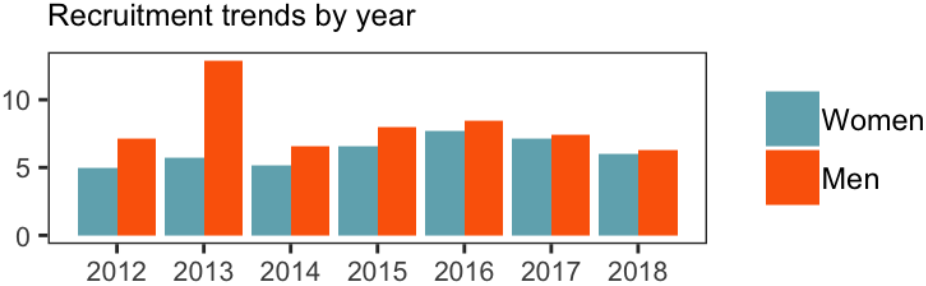
Recruitment of Men vs. Women by year, between 2012 and 2018.

### Recommendations for improved support of new principal investigators

The career trajectory of an independent academic researcher is diverse and these data support that there are many different variations of routes to be taken. However, despite the individual nature of each academic appointment, these data do highlight a few overarching issues affecting many of our respondents. We therefore make the following practical suggestions for changes in policy that could be adopted by both host institutions and funders, to ensure that the potential of all newly recruited independent investigators is fully maximised.

### Actions by host institutions

- Ensure that all research fellows receive a formal annual appraisal.
- Ensure that all new independent researchers are appointed a departmental mentor.
- Arrange that all research fellows be assessed for a proleptic appointment and support for senior fellowship applications within 24 months of the end of their career development fellowship (year 3/4).
- Make assessment criteria for promotion/proleptic appointments transparent and ensure that these criteria are communicated to the fellow or lecturer from the start of their appointment.
- Research fellows should be permitted to spend the majority of their time on research activities and not expected to pick up significant teaching load.
- Consider a standard policy that newly appointed lecturers should be appointed on grade 8 and considered for promotion to senior lecturer (grade 9) upon successfully winning their first major research grant.
- Consider a standard policy that research fellows should be appointed at grade 9 if they start their position with a major external research grant.
- Consider a standard policy that university-funded fellows should be appointed at grade 8 and considered for promotion to grade 9 upon successful application for a major external research grant.
- Limit the number of graduate research project and master’s project students for new principal investigators, the number not to exceed the total number of staff in the research group (Postdoctoral researchers, PhD students, research assistants, and PI).
- Give priority to new PIs when selecting PhD projects for university administered doctoral training programmes and award a proportion of PhD studentships to new PIs.

### Actions by funders

- Reconsider the decision to fund PhD studentship primarily through large university administered training programs. Understanding that this approach mostly benefits more senior labs.
- Consider offering a PhD studentship as part of a career development fellowship or new investigator award.
- Funders should withhold funds from host institutions if commitments such as facility access and lab space is not provided as specified and committed to in the application.
- Funders should engage with host institutions to monitor the career progression of research fellows, to ensure equal and fair assessment of fellows and lecturers.
- Funders should monitor and consider diversifying their funding by awarding grants and fellowships to non-Russell group universities and institutions outside of London.
- Consider a standard policy to recommend that research fellows should be appointed at grade 9, or the equivalent of a senior lecturer.

### Advice when applying for independent positions

When making the transition and applying for your first independent position, take control. Know what you will be provided with by your host, and your funder. Don’t leave arrangements to chance.

- Be aware that you are recruited to become part of a department, you should fully understand the department’s goals and what role you are expected to fill as you join.
- Talk details. Ask to see the lab space you will be working in, ask who will provide lab basics like the fridges and freezers, find out what administrative support you will have (ordering, finance, travel bookings)
- Simple and obvious, make sure that your starting grade/salary is set as suggested above (see actions by host institutions). New lecturers at grade 8 moving to grade 9 after being awarded the first major grant. Fellows starting with a major research grant should start on grade 9 or senior lecturer equivalent. Where you start in the system will impact your future promotions.
- Ask to speak with other new PIs, either in your department or within your institution to find out how the host institution works and how they were recruited and supported.
- Make sure you have a mentor in your department.
- Make sure you have an annual review, and a time frame for review for a promotion or proleptic appointment.
- Find your peers, and talk with them often, other new PIs will be your best support network. Starting a lab can be a lonely business and very different from being a postdoc.
- Don’t assume that the person you are negotiating with has the power or authority to agree to what you are requesting, be aware that heads of department can change…
- And so finally, if you are promised certain support from your host, or need access to particular equipment for example…**Get. Everything. In. Writing**.

## Methods and Statistical analysis

The survey was conducted using convenience sampling, with most participants finding out about it through Twitter or forwarded email invitations. The majority of those responding were in the life sciences, in part because of the networks that the survey was circulated around, and partly due to the language used in the survey (‘PI’ does not have the same meaning in the social sciences, for instance). We do not claim that this is a fully representative sample. However, we do feel that it allows us to say important things about at least a significant subgroup of New PIs in the UK.

Much of the analysis consists of simple descriptive statistics - that is, looking at the distribution of individual variables. However, where we were interested in the relationship between variables, we used a mix of ordinal logit regressions and chi-squared tests depending on the nature of the relationship being studied. Ordinal regression allowed us to control for multiple factors, to be more sure that the relationship that we found was not a result of (at least measured) confounding factors. Full details of these models can be found in the supplementary data, with some statistics indicating significance reported in the text.

## Supporting information

Statistical calculations

## ACKNOWLEDGEMENTS

We thank all the members of UK_NewPI slack group for active discussion, honesty and productive practical suggestions for improvement which could be made to ease this difficult career stage. Thank you also to Dr Yanlan Mao (MRC LMCB) and Dr Ian Sudbery (University of Sheffield) for input on the structure and content of the survey questions, and to the MRC LMCB Athena Swan and EDIC committee for their discussions and engagement in this project. Thank you the #eLIFEAmbassadors who have supported this project and thank you to the funders and universities who have engaged with us throughout the analysis and are enthusiastic to listen and to make change.

**Dr. Sophie E. Acton** is a Cancer Research UK career development fellow at University College London | **Dr. Andrew J.D. Bell** is a lecturer at the University of Sheffield | Dr Christopher P. Toseland is an MRC Career Development awardee at University of Kent | **Dr. Alison Twelvetrees** is a a Vice Chancellor’s fellow at the University of Sheffield.

